# Cross-regulation of amino acid synthesis and catabolic electron transfer in bacteria

**DOI:** 10.1101/2024.12.27.630285

**Authors:** Hisae Mogi, Keisuke Tomita, Atsumi Hirose, Erika Yoshino, Takuya Kasai, Atsushi Kouzuma, Kazuya Watanabe

## Abstract

Amino acids are essential for life, serving primarily as the building blocks of proteins. Although the intracellular metabolic fate of amino acids is known, their roles in the global regulation of metabolic pathways are poorly characterized. Here, we investigated the function of amino acids as signaling molecules that modulate the catabolic activity of bacterial cells. Using the model bacterium *Shewanella oneidensis* MR-1, we showed that methionine activates catabolic processes, including extracellular electron transfer, at submillimolar concentrations. This regulation is mediated by MetR, a transcription factor that regulates methionine biosynthesis. In addition, the MetR-mediated regulation of catabolic electron transfer was observed in *Aeromonas hydrophila*. These findings reveal methionine-induced signaling cascades that cross-regulate amino acid biosynthesis and catabolism in bacteria. Our study highlights a novel physiological role for amino acids, suggesting their involvement in the coordinated regulation of anabolism and catabolism, particularly in bacteria that thrive in nutrient-limited habitats.

## Introduction

Balancing anabolic and catabolic metabolism is essential for the survival of organisms, especially microorganisms that inhabit nutrient-poor or energy-limited environments^1,2^. These microorganisms must efficiently utilize the available nutrients in their environment and allocate the energy obtained from catabolism (e.g., respiration) to the synthesis of cellular components (e.g., amino acids and proteins). Studies have revealed the physiological functions of specific regulators that individually control the expression of anabolic and catabolic genes in microbial cells^3^. However, little information is available on the interactive regulation of anabolic and catabolic metabolism.

*Shewanella oneidensis* MR-1 is a facultative anaerobic bacterium belonging to the class Gammaproteobacteria^4^. *S. oneidensis* MR-1 can utilize various anaerobic electron acceptors, such as fumarate, nitrate, dimethyl sulfoxide (DMSO), trimethylamine *N*-oxide (TMAO), and metal oxides^5^. Due to its ease of culture and genetic manipulation, as well as its diverse respiratory activities, *S. oneidensis* MR-1 is used as a model organism to study how bacteria adapt to and thrive in redox-stratified environments with limited nutrient and energy availability^6^. Moreover, MR-1 is extensively studied as a host for microbial electrochemical technologies (METs), such as microbial fuel cells and electro-fermentation systems, because it can link intracellular metabolism (redox reactions) to electrodes through the extracellular electron transfer (EET) pathway^7,8^. Understanding how MR-1 regulates anaerobic respiration in response to environmental cues is critical for advancing our knowledge of microbial survival strategies and optimizing its applications in METs.

MR-1 uses the cyclic AMP receptor protein (CRP) to activate the expression of many anaerobic respiratory genes, including those involved in the reduction of metal oxides and electrodes (*omcA* and *mtrCAB*) and DMSO (*dmsEFABGH*)^9,10^. CRP binds directly to the upstream regions of *omcA* and *mtrCAB* and activates their transcription upon receiving the intracellular second messenger cAMP^10^. However, the specific environmental stimuli that trigger the expression of CRP-regulated genes are unidentified, warranting further investigation to elucidate the mechanisms that regulate the anaerobic respiration of MR-1. On the other hand, our previous study revealed that MR-1 downregulates the expression of amino acid biosynthesis pathway components under anaerobic, electron acceptor-limited conditions, thus requiring an external supply of amino acids for fermentative growth in defined media lacking terminal electron acceptors^11^. This finding suggests the presence of unknown regulatory crosstalk between amino acid biosynthesis and anaerobic respiration pathways in MR-1.

Many bacteria, including *S. oneidensis* MR-1, possess intrinsic pathways for amino acid biosynthesis. These pathways are regulated by specific transcription factors that respond to the intracellular concentrations of individual amino acids^3^. Through such regulatory mechanisms, de novo amino acid synthesis is suppressed as intracellular amino acid levels rise. The synthesized amino acids are directed to protein synthesis, including the production of catabolic enzymes. However, the influence of intracellular amino acid levels on the catabolic activity of bacterial cells remains poorly understood. In this study, we hypothesize that amino acid availability affects the catabolic (respiratory) activity of MR-1 through an unknown regulatory mechanism. Given that MR-1 can use an electrode as an electron acceptor for respiration, its respiratory activity (i.e., electron transfer rate) can be readily quantified in real-time by monitoring current generation in an electrochemical cell (EC)^12^. Therefore, we investigated the effect of external amino acid supplementation in defined media on current generation by MR-1 to test our hypothesis.

## Results

### Effects of amino acids on the EET activity of MR-1

To explore the amino acids that influence the catabolic electron transfer activity of MR-1, we assessed current generation by this strain in ECs filled with lactate minimal medium (LMM, containing 10 mM lactate as the carbon and energy source) supplemented with one of the 20 amino acids (130 µM each) (Supplementary Fig. S1). The initial screening assays, followed by confirmatory measurements (Fig. 1a, b), revealed that methionine (Met) supplementation increased current generation 1.5-fold. To investigate the dose-dependent effect of Met supplementation, current generation was evaluated in the presence of 0. 013 and 1.3 µM Met (Supplementary Fig. S2). Current generation increased significantly at 1.3 µM Met, with a positive correlation between increasing Met concentrations and current generation. However, supplementation with 130 µM Met did not affect the growth of MR-1 in LMM under either aerobic or anaerobic fumarate-reducing conditions (Supplementary Fig. S3). These observations suggest that Met, or its intracellular metabolite, acts as a signaling molecule that modulates the EET activity of MR-1, rather than serving as a growth-promoting factor in minimal media. Moreover, the addition of homocysteine (hCys), a metabolic intermediate in the Met biosynthesis pathway^13^, negatively affected current generation by MR-1 (Supplementary Fig. S1, Fig. 1a, b). This finding further supports the hypothesis that Met metabolism is related to the regulation of the EET activity of MR-1 cells.

**Fig. 1.**
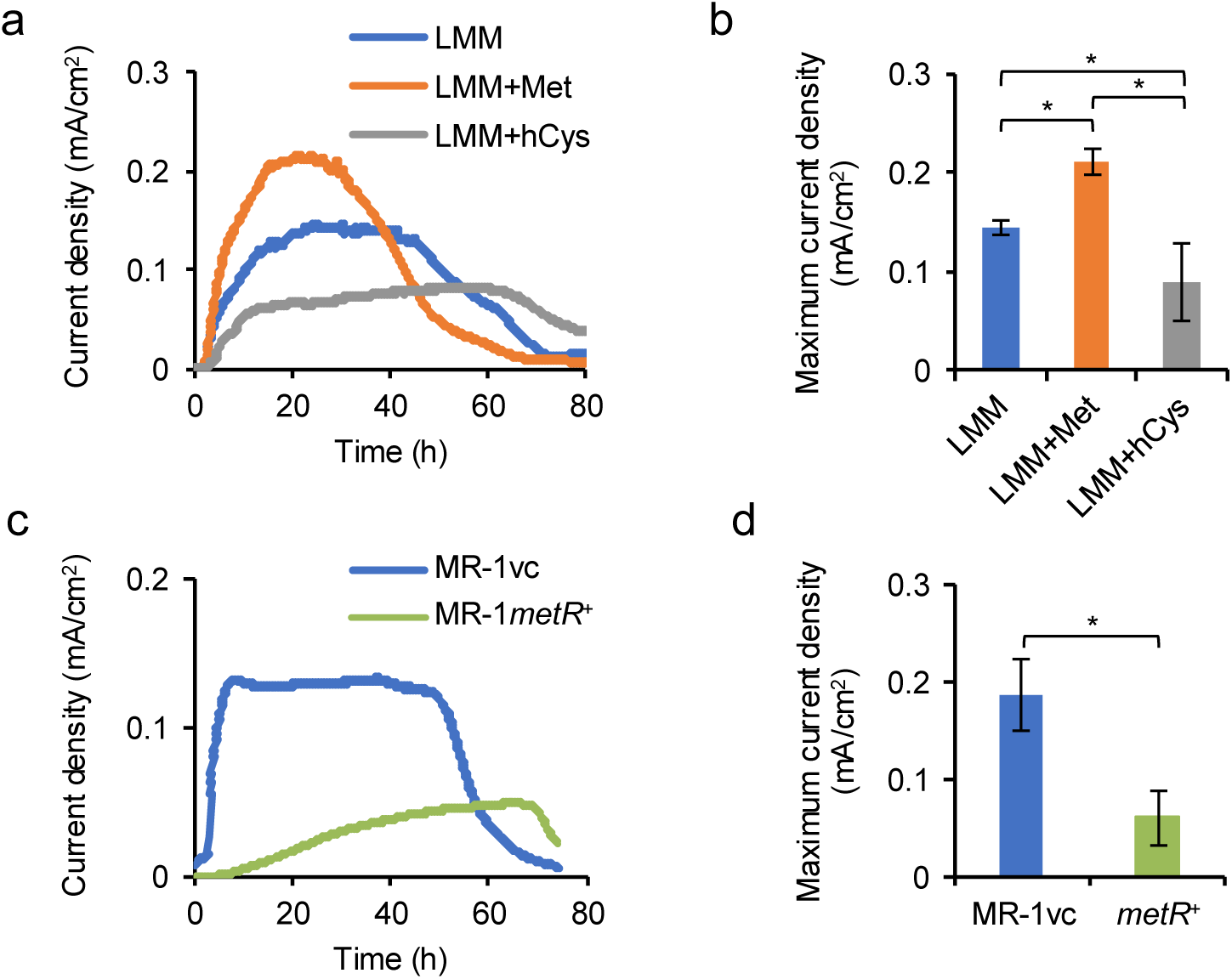
Effects of Met, hCys, and *metR* expression on current generation by *Shewanella oneidensis* MR-1. (a, b) Representative current density versus time curves (a) and maximum current densities (b) for wild-type MR-1 in ECs containing LMM supplemented with 130 µM Met or hCys. (c, d) Representative current density versus time curves (c) and maximum current densities (d) for MR-1vc and MR-1*metR^+^* in ECs containing LMM. In panels b and d, the bars and error bars represent the means and standard deviations, respectively (*n* = 3 biological replicates). Asterisks indicate statistically significant differences (*P* < 0.05; one-way ANOVA followed by HSD test in panel B, Student’s *t* test in panel d).

### Effects of Met supplementation on the transcriptomic profile of MR-1

To investigate the impact of Met supplementation on the expression of EET-related and other catabolic genes in MR-1, we conducted a comparative transcriptomic analysis of MR-1 cells cultured in LMM with or without 130 µM Met. Total RNA was extracted from aerobically cultured MR-1 cells to minimize the potential confounding effects associated with changes in anaerobic respiratory activity. Out of 4,214 genes in the MR-1 genome, 599 were upregulated and 555 were downregulated in the presence of 130 µM Met, with statistical significance at *P* <0.05 and a log_2_-fold change (log_2_ FC) threshold of ≥1 or ≤–1 (Supplementary Data S1, S2). The genes involved in Met biosynthesis (*met* genes) were most significantly downregulated under Met-supplemented conditions (Table 1 and Supplementary Data S2), likely due to a feedback regulatory mechanism for Met biosynthesis^14^. Further analysis focused on genes related to EET and anaerobic respiration, revealing consistent upregulation of genes encoding EET components (*cymA*, *omcA*-*mtrCAB*, and *fccA*) and DMSO reductase (*dms* genes) in response to Met supplementation (Table 1, Supplementary Data S1). The upregulation of *cymA* is noteworthy, because CymA, the inner membrane-anchored periplasmic cytochrome *c* essential for EET, functions as a critical electron transfer hub in many anaerobic respiratory pathways, including those for fumarate, DMSO, and nitrate reduction^15^. In addition, Met supplementation upregulated the *hem* and *ccm* genes, which are involved in heme synthesis^16^ and cytochrome *c* maturation^17^, respectively (Table 1, Supplementary Data S1). Collectively, these findings suggest the existence of a regulatory mechanism that coordinately activates the expression of genes involved in anaerobic electron transfer in response to Met availability.

**Table 1.**
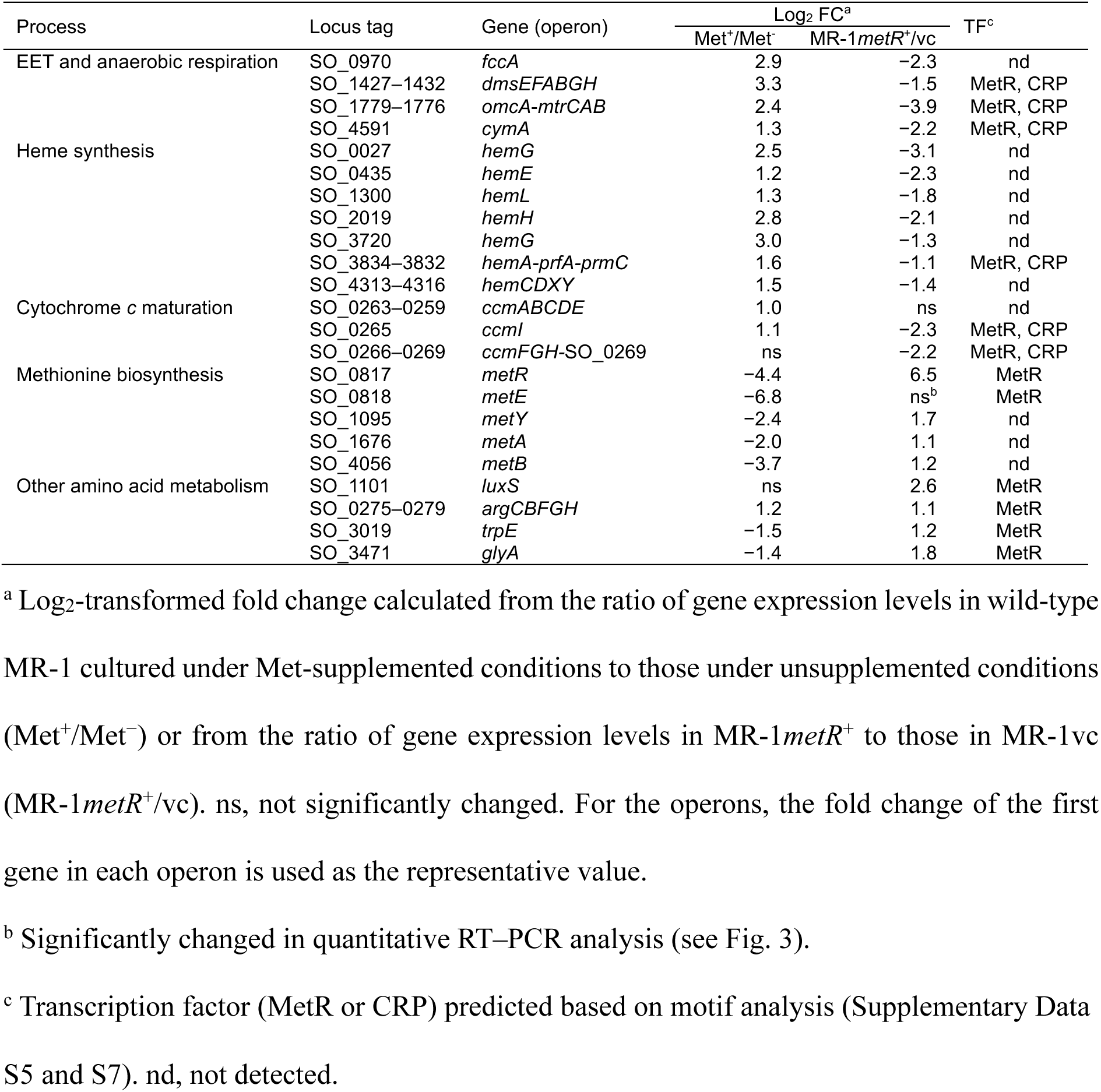
List of selected Met-responsive or MetR-regulated genes in MR-1.

### Regulation of anaerobic electron transfer genes by MetR

The MR-1 genome encodes an ortholog of MetR, a transcriptional activator of *met* genes, which shares 46% amino acid identity with the well-characterized *Escherichia coli* MetR^18^. We hypothesize that MetR negatively regulates the transcription of genes involved in EET and other anaerobic respiration pathways, given that current generation was enhanced by Met and suppressed by hCys, an inducer of MetR^19^ (Fig. 1a, b), and that Met supplementation significantly downregulated *metR* transcription (Table 1). To test this hypothesis, we constructed a *metR* deletion strain (Δ*metR*), a complemented strain (Δ*metR*-C), and an overexpression strain (MR-1*metR*^+^) and characterized the physiological properties of these MR-1 derivatives. Growth assays in LMM (Supplementary Fig. S4) and lysogeny broth (LB) medium (Supplementary Fig. S5) revealed that the *metR* deletion strain (transformed with the control vector pBBR1MCS-5; Δ*metR*vc) exhibited a severe growth defect in the minimal medium, which was rescued in Δ*metR*-C. Given this growth defect of Δ*metR*, we focused subsequent analyses on characterizing MR-1*metR*^+^. Measurements of current generation in ECs (Fig. 1c, d) demonstrated that MR-1*metR*^+^ produced significantly lower current than the vector control strain (MR-1vc; harboring pBBR1MCS-5).

Additionally, we examined the growth of MR-1*metR*^+^ in the presence of different electron acceptors using LB (amino acid-rich) medium to minimize the influence of *metR* expression on amino acid metabolism and the associated cell growth. MR-1*metR*^+^ exhibited markedly impaired growth under fumarate-, DMSO-, and nitrate-reducing conditions (Fig. 2), while showing comparable growth to MR-1vc under aerobic conditions (Supplementary Fig. S6) and anaerobic TMAO-reducing conditions (Fig. 2). These findings indicate that MR-1*metR*^+^ has a diminished capacity for EET and fumarate, DMSO, and nitrate respiration.

**Fig. 2.**
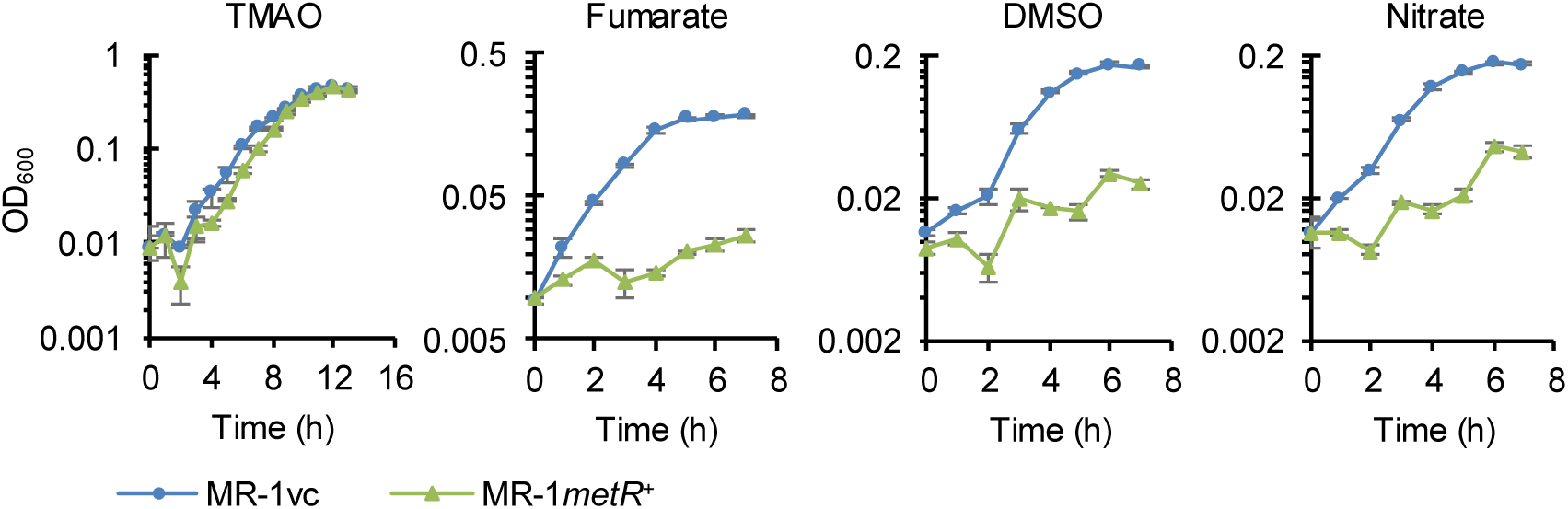
Growth of MR-1vc and MR-1*metR^+^* under anaerobic respiration conditions. *Shewanella oneidensis* was cultured in LB medium supplemented with TMAO, fumarate, DMSO, or nitrate as the electron acceptor. Bars and error bars indicate means and standard deviations, respectively (*n* = 3 biological replicates).

To investigate the mechanism of impaired current generation and anaerobic growth observed in MR-1*metR*^+^, we conducted a comparative transcriptomic analysis of MR-1*metR*^+^ and MR-1vc. Both strains were anaerobically cultured in LMM containing TMAO, an electron acceptor that MR-1*metR*^+^ can utilize (growth curves are shown in Supplementary Fig. S4). The overexpression of *metR* led to significant transcriptional changes in 722 genes, with 308 genes upregulated and 414 downregulated (Supplementary Data S3, S4). Notably, many differentially expressed genes in MR-1*metR*^+^ exhibited expression patterns opposite to those observed with Met supplementation. Specifically, many genes involved in EET and anaerobic respiration, as well as several *hem* (heme synthesis) and *ccm* (cytochrome *c* maturation) genes, were downregulated in MR-1*metR*^+^, whereas *met* genes were largely upregulated (Table 1). This trend was further confirmed by quantitative RT-PCR analysis for four selected genes (*omcA*, *fccA*, *dmsB*, and *metE*) (Fig. 3). These findings suggest that the impaired current production and anaerobic growth observed in MR-1*metR*^+^ are related to the transcriptional repression of genes involved in the EET and anaerobic respiratory pathways.

**Fig. 3.**
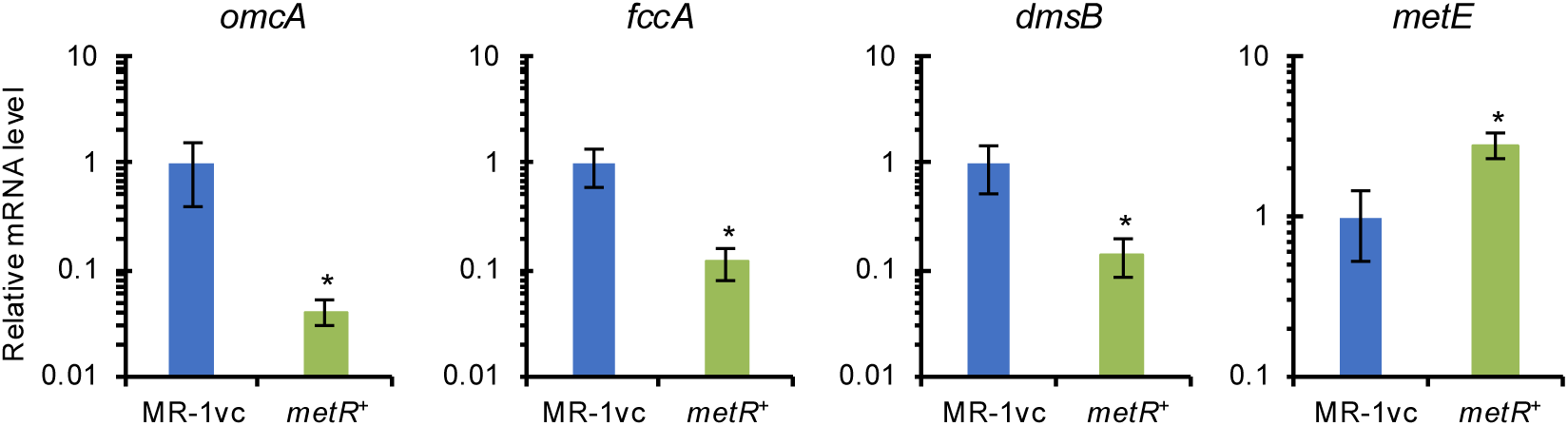
Differential expression of anaerobic respiration and methionine biosynthesis genes in MR-1*metR^+^*. The expression levels of *omcA*, *fccA*, *dmsB*, and *metE* were analyzed by quantitative RT-PCR using total RNA extracted from cells cultured anaerobically in LMM supplemented with TMAO as the electron acceptor. Results are presented as the relative levels of mRNA expression in MR-1vc cells. Bars and error bars represent means and standard deviations, respectively (*n* = 3 or 4 biological replicates). Asterisks indicate statistically significant differences (*P* < 0.05; Student’s *t* test).

### Binding of MetR to the upstream regions of anaerobic electron transfer genes

We performed electrophoretic mobility shift assays (EMSA) to investigate the direct regulation of EET and anaerobic respiratory genes by MetR (Fig. 4). A 13-bp palindromic consensus sequence for MetR binding (5′-TGAANNNNNTTCA-3′) has been reported in *E. coli*, *Salmonella typhimurium*, and *Vibrio harveyi*^20,21^, and similar sequences were found in the upstream region of *metR* (the *metR*–*metE* intergenic region) as well as in the upstream regions of several differentially expressed genes in MR-1*metR*^+^, including *luxS*, *glyA*, *omcA*, *cymA*, and *dmsE*. Based on the upstream regions of these six genes, we generated 40-bp DNA probes containing the putative 13-bp MetR-binding sequences (Fig. 4a) and examined the binding of purified MetR (Fig. 4b) to these probes. When MetR was incubated with the *omcA* probe, shifted bands were observed in a protein concentration-dependent manner, whereas the shift disappeared in the presence of a specific DNA competitor (excess unlabeled *omcA* probe) (Fig. 4c). Similar shifts were observed when MetR was incubated with the other five DNA probes, whereas no shift was detected with the mutated *omcA* probe, in which the palindromic nucleotides in the 13-bp consensus sequence had been replaced (Fig. 4d). These results demonstrate that MetR directly binds to the upstream regions of the six tested genes containing the predicted MetR-binding sequences.

**Fig. 4.**
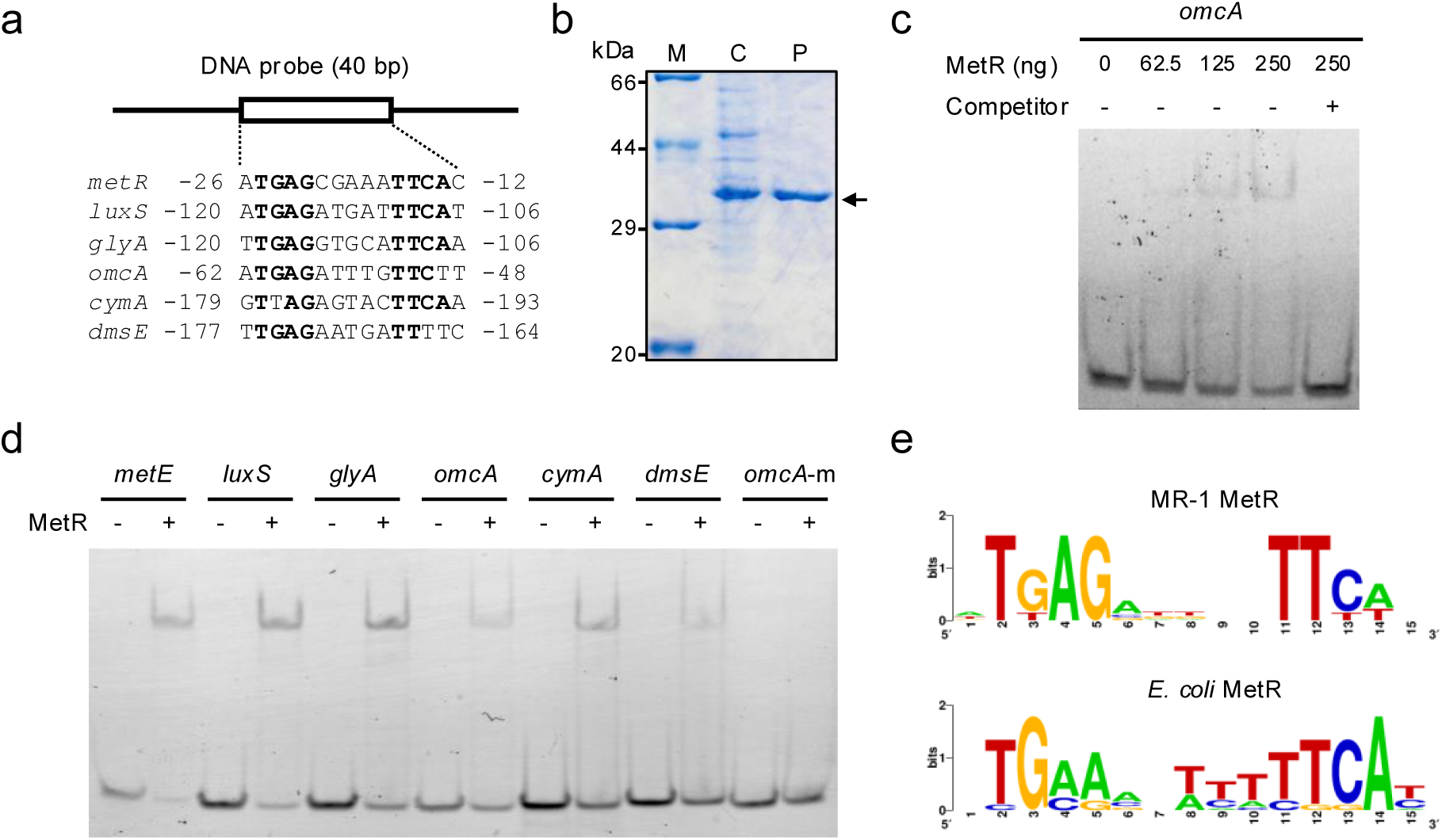
Binding of MetR to DNA regions upstream of the selected MetR-regulated genes. (a) DNA probes used in EMSA. Nucleotides commonly found in the MetR-binding sequences are shown in bold. Numbers on either side of each DNA sequence indicate positions relative to the start codon (+1) of the downstream gene. (b) SDS-PAGE of purified MetR. The arrow marks the band corresponding to C-his-MetR (33.4 kDa). Lane M, molecular weight marker; Lane C, *E. coli* BL21(DE3)(pET-metR) crude extract; and Lane P, purified extract. (c) EMSA using MetR and *omcA* probe. The Cy-3-labeled probe was incubated with 0–250 ng MetR in the presence (+) or absence (–) of a specific competitor (a 5000-fold excess of unlabeled *omcA* probe). (d) EMSA using DNA probes of selected genes involved in anaerobic respiration and methionine metabolism. Labeled probes were incubated in the presence (+) or absence (–) of 500 ng of MetR. The mutated *omcA* probe (*omcA*-m), in which the palindromic eight nucleotides in the 13-bp MetR-binding consensus sequence were substituted with guanine residues, was used as a negative control. (e) Site-specific nucleotide frequencies in the MetR-binding DNA sequences. The sequence logo representing the MetR-binding sequences in MR-1 was generated using the six sequences shown in panel a. For comparison, the sequence logo for known MetR-binding sequences in *E. coli* K-12 is shown. The height of each letter represents the degree of sequence conservation measured in bits.

### Prediction of the MetR regulon in MR-1

Based on the six MetR-binding sequences identified by EMSA (Fig. 4a), we constructed a position-specific scoring matrix (PSSM) for the consensus MetR-binding sequence in MR-1 and visualized it as a sequence logo (Fig. 4e). The 13-bp MR-1 MetR-binding sequence (consensus: 5′-TGAGNNNNNTTCA-3′) was largely identical to that of *E. coli*, although the substitution of the fourth nucleotide from A to G slightly disturbed palindromic symmetry. Using PSSM, we scanned the MR-1 genome (500 bp upstream regions of each gene) and identified putative MetR-binding sites in the upstream regions of 151 genes (including the six genes used for constructing PSSM; Supplementary Data S5). Of these, 37 genes were differentially expressed in MR-1*metR*^+^, with 19 upregulated and 18 downregulated (Supplementary Data S5). These analyses suggest that these 37 genes, along with their associated operons, are directly regulated by MetR. Based on this information and the cotranscription patterns of the gene clusters observed in the transcriptome analysis (Supplementary Data S3, S4), we predicted the MetR regulon in the MR-1 genome (Supplementary Data S6). Notably, the MetR regulon included several *hem* and *ccm* genes, as well as *cymA*, *omcA*-*mtrCAB*, and *dmsEFABGH*, which were consistently downregulated in MR-1*metR*^+^ (Supplementary Data S6). Genes specifically involved in TMAO reduction (*torECAD*)^22,23^ are not included in the MetR regulon, consistent with the observation that MR-1*metR*^+^ can grow under TMAO-reducing conditions (Fig. 2). The analysis predicted that MetR directly activates the transcription of genes involved in the metabolism of arginine (*argCBFGH*), phenylalanine (*pheA*), tryptophan (*trpE*), and glycine (*glyA*), as well as methionine (*metE* and *luxS*). These results suggest that MetR functions as a global regulator, repressing the transcription of many genes involved in anaerobic electron transfer and activating the transcription of some genes involved in amino acid metabolism.

MR-1 uses CRP to activate the transcription of genes involved in EET and other anaerobic respiratory systems, excluding TMAO reduction^9^. To explore the overlap between the MetR and CRP regulon in this strain, we scanned the MR-1 genome using PSSM for the CRP-binding sequences identified in *E. coli*, given that MR-1 CRP shares high amino acid identity (89%) with *E. coli* CRP. This analysis identified putative CRP-binding sites in the upstream regions of 369 genes (Supplementary Data S7), 12 of which, including *omcA*, *dmsE*, *cymA*, and *ccmF*, overlapped with the MetR regulon (Table 1, Supplementary Data S6). Notably, the CRP-binding site upstream of *omcA*, which was experimentally confirmed in our previous study^10^, is located upstream of the MetR-binding site (Supplementary Fig. S7). Similar arrangements of CRP– and MetR-binding motifs are also present in the upstream regions of *cymA*, *dmsE*, and *ccmF* (Supplementary Data S6). These arrangements, together with the observed downregulation of these four genes (and associated operons) in MR-1*metR*^+^ (Table 1, Fig. 3), suggest that MetR represses their transcription by interfering with CRP-dependent transcriptional activation.

### Inhibition of EET by *metR* expression in *Aeromonas hydrophila*

Given that MetR is a transcription factor widely conserved across proteobacteria^14^, we hypothesize that the MetR-dependent repression of anaerobic electron transfer systems in MR-1 is conserved in other bacteria, particularly those closely related to *Shewanella*. To test this hypothesis, we engineered *Aeromonas hydrophila* ATCC 7966, a bacterium with EET activity^24^ and relatively close phylogenetic proximity to *Shewanella*^25^, to overexpress its native *metR* (AHA_1955). The resulting strain, designated AH*metR*^+^, exhibited similar growth to the vector control strain (AHvc) when cultured in LMM under aerobic and fumarate-reducing conditions (Fig. 5a). However, AH*metR*^+^ generated lower current in EC compared with AHvc (Fig. 5b), demonstrating that *metR* expression negatively impacts EET in *A. hydrophila*. For comparison, we constructed an *E. coli* strain (EC*metR*^+^) that overexpressed its native *metR* and assessed its growth under anaerobic respiration conditions using fumarate and nitrate as terminal electron acceptors. However, the growth of EC*metR*^+^ was comparable to that of the vector control strain (ECvc) under these conditions (Supplementary Fig. S8), suggesting that in *E. coli*, MetR does not play a role in regulating fumarate and nitrate respiration.

**Fig. 5.**
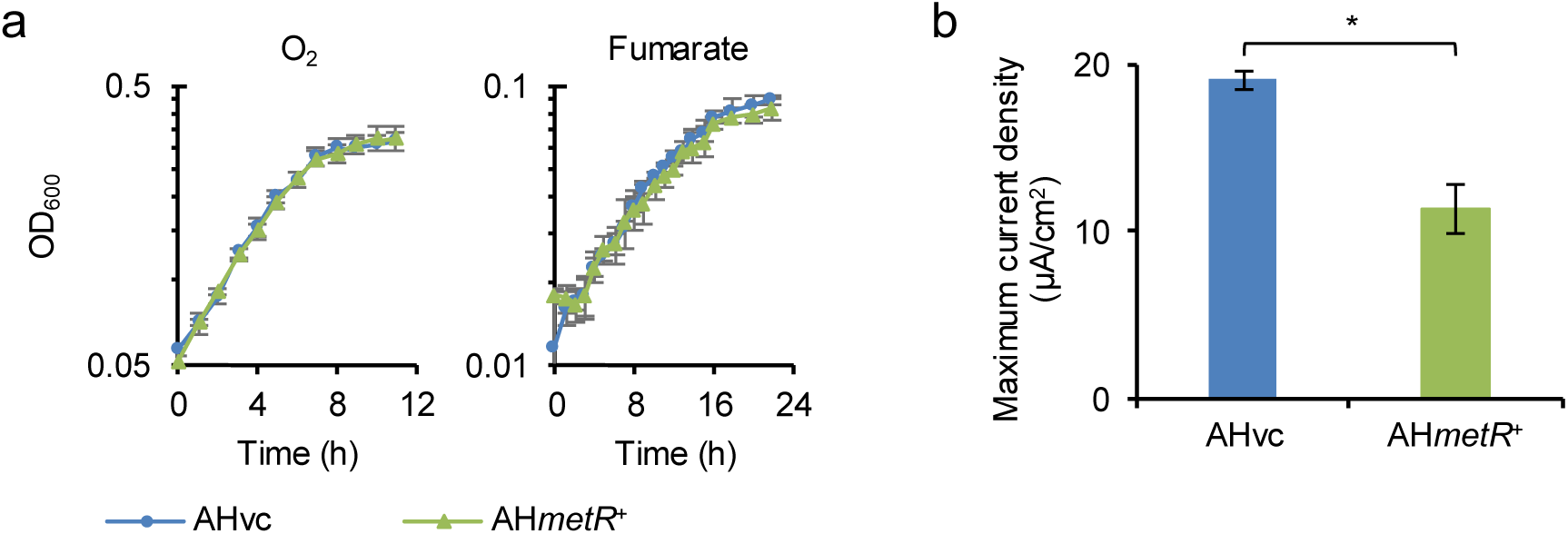
Effect of *metR* expression on the respiratory activity *of Aeromonas hydrophila*. (a) Growth of AHvc and AH*metR^+^*in LMM under aerobic (O_2_-reducing) and fumarate-reducing conditions. (b) Maximum current densities in ECs inoculated with AHvc and AH*metR^+^.* In all panels, error bars represent standard deviations (*n* = 3 biological replicates). Asterisks indicate statistically significant differences (*P* < 0.05; Student’s *t* test).

To predict MetR-regulated genes in *A. hydrophila*, we scanned its genome using PSSM for the MetR-binding sequences identified in MR-1 (Supplementary Data S8). We identified a putative MetR-binding sequence in the upstream region of *ccmF*, suggesting that *A. hydrophila* uses MetR to negatively regulate cytochrome *c* maturation. However, no putative MetR-binding sequences were found in the upstream regions of the known EET component genes *mtrCAB* (AHA_2764–2766), *pdsA* (AHA_2763), and *netBCD* (AHA_2762-2760)^24^. These results, together with the comparatively weaker repression of EET in *A. hydrophila* relative to that observed in MR-1 (Fig. 1, 5b), suggest that the influence of MetR on the repression of anaerobic electron transfer systems in *A. hydrophila* is more limited than its effects in MR-1.

## Discussion

In *E. coli* and *Salmonella*, MetR activates the transcription of *metE*, which encodes the cobalamin-independent methionine synthase, in the presence of hCys as an inducer^19,20^. In these enteric bacteria, *metR* and *metE* are located adjacently in opposite orientations, with overlapping promoter regions. Consequently, MetR activates *metE* transcription while repressing its own expression. In addition to this autoregulatory mechanism, *metR* transcription is repressed by MetJ and its corepressor, *S*-adenosylmethionine (SAM)^13^.

Synthesized from methionine, SAM enhances the DNA-binding activity of MetJ, thereby facilitating the transcriptional repression of MetJ-regulated genes, including *metR* and other Met biosynthesis (*met*) genes, such as *metA*, *metB*, *metC*, *metE*, and *metF*^26^. These regulatory systems ensure that *met* expression is upregulated when intracellular Met levels are low. In MR-1, *metR* and *metE* are similarly arranged in opposite directions, and the intergenic region contains a MetR-binding sequence (Fig. 4). The deletion of *metR* in this strain led to growth inhibition in minimal medium (Supplementary Fig. S4), consistent with observations in *E. coli*^27^. Together with the transcriptomic profiles observed in the presence of Met or in MR-1*metR*^+^ (Table 1), these findings suggest that MR-1 employs regulatory systems for *met* genes analogous to those conserved in enteric bacteria.

In this study, we demonstrated that MR-1 utilizes MetR not only to activate the Met biosynthesis pathway but also to repress anaerobic electron transfer systems. The regulatory functions of MetR proposed are summarized in Fig. 6. Under Met-limited conditions, *metR* expression is upregulated, leading to the transcriptional activation of *metE* and several other amino acid biosynthesis genes, whereas the expression of genes encoding cytochrome *c* proteins involved in anaerobic respiration (*omcA*, *cymA*, and *dms* operon) and several heme synthesis (*hem*) and cytochrome *c* maturation (*ccm*) genes is repressed (Fig. 6a). Given that hCys inhibited the EET activity of MR-1 (Fig. 1a, b), it is likely that hCys acts as a corepressor in the MetR-dependent repression of these genes. Conversely, under Met-rich conditions, *metR* expression is downregulated, leading to the transcriptional activation (derepression) of anaerobic electron transfer genes (Fig. 6b).

**Fig. 6.**
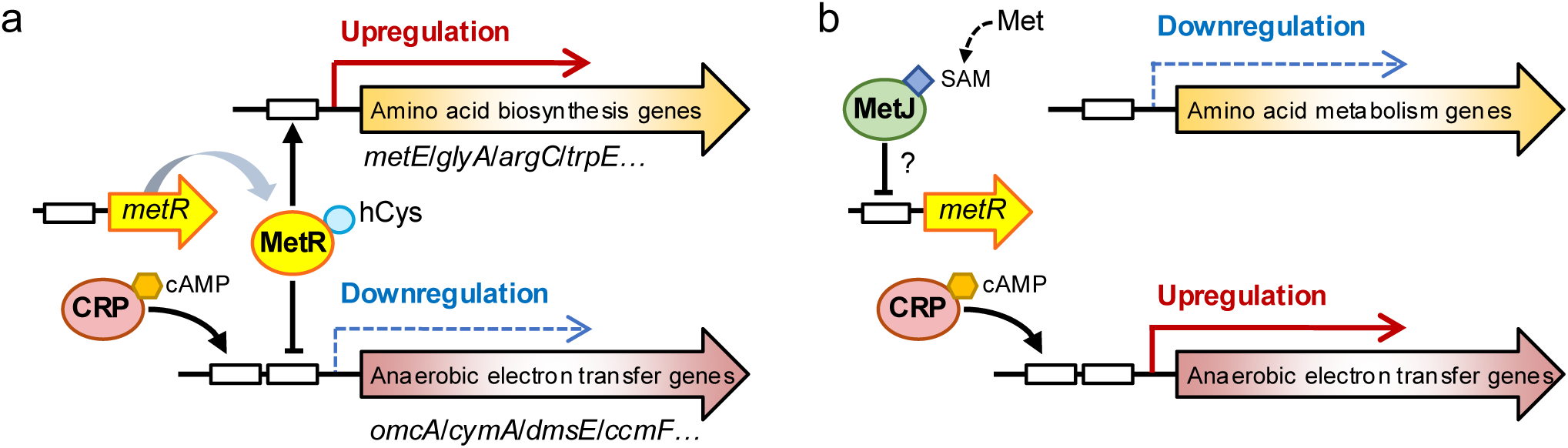
Proposed mechanisms of MetR-mediated regulation of anaerobic electron transfer and amino acid biosynthesis pathways. (a) Regulation under Met-limited conditions. (b) Regulation under Met-rich conditions. Boxes represent upstream regulatory regions bound by transcription factors.

This regulatory mechanism may represent a survival strategy that bacteria have evolved to optimize growth in response to nutrient and energy availability. Met is the most energetically expensive amino acid to synthesize under anaerobic conditions^28^. However, anaerobic respiratory bacteria, particularly those expressing EET pathways, must expend considerable energy to synthesize heme and cytochrome *c* for respiration^29,30^, despite the limited energy resources available in their environment. Thus, when these bacteria have access to externally supplied Met, it is reasonable that they conserve energy by repressing their own Met biosynthesis pathways, allowing them to allocate more energy to the production of cytochrome *c* proteins required for anaerobic respiration. Conversely, when Met is scarce, the bacteria optimize their intracellular energy allocation by downregulating anaerobic electron transfer and upregulating Met biosynthesis. We propose that certain bacteria, including *S. oneidensis*, use intracellular Met levels as an indicator of overall amino acid levels, likely due to the energetic difficulty of its biosynthesis, thereby allowing efficient expression of anaerobic electron transfer pathways in response to intracellular amino acid and energy levels.

The MetR regulon in MR-1, predicted based on the MetR-binding sequences identified by EMSA (Fig. 4) and the transcriptomic profile of MR-1*metR*^+^, includes *metE* (*metR*), *luxS*, and *glyA*, as conserved in other proteobacteria^14^. However, the MetR regulon in this strain is distinct in that it also includes *arg* and *trpE*, suggesting that MR-1 employs MetR to regulate a relatively large number of amino acid biosynthesis pathways. Notably, our analysis suggests that a part of the MR-1 MetR regulon, including several genes involved in anaerobic respiration (*cymA*, *omcA*, and *dmsE*) and cytochrome *c* maturation (*ccmI* and *ccmF*), overlaps with the CRP regulon (Table 1 and Supplementary Data S6). Based on our data and current knowledge of CRP function in MR-1^9,10^, it appears that many of the MetR– and CRP-regulated genes are transcriptionally upregulated by CRP and downregulated by MetR. A similar competitive regulatory mechanism by CRP and MetR was reported in *V*. *harveyi*^21^, where both regulators, with the quorum-sensing regulator LuxR, bind to the *lux* promoter, thereby modulating bioluminescence in response to nutrient availability and cell density. Analogously, the regulation of anaerobic electron transfer systems by CRP and MetR in MR-1 may enable flexible control of respiratory activity in response to fluctuations in nutrient and energy availability.

MetR is involved in the regulation of virulence in *Vibrio cholerae*^31^, secondary metabolite synthesis in *Serratia marcescens*^32^, and swarming in *Pseudomonas aeruginosa*^33^. These observations and our results indicate that MetR plays distinct physiological roles in phylogenetically diverse bacteria. Our data further suggest that MetR is involved in the regulation of EET in *A. hydrophila* (Fig. 5b). However, predictions of MetR-binding sequences suggest that in this bacterium, MetR does not directly regulate genes encoding cytochrome *c* involved in the EET pathway, but rather controls several *ccm* genes (Supplementary Data S8). Cytochrome *c* maturation likely contributes to the overall activity of respiratory electron transfer systems under both aerobic and anaerobic conditions.

However, in MR-1, the expression of *ccm* genes is upregulated under oxygen-limited conditions, in coordination with genes involved in anaerobic respiration^34^, indicating the particular significance of cytochrome *c* maturation in anaerobic respiration. Because EET requires a substantial number of cytochrome *c* proteins^29,30^, it is plausible that the *A. hydrophila metR*-overexpressing strain (AH*metR*^+^) exhibited partially impaired current production due to the repression of cytochrome *c* maturation, despite showing no growth defects under either aerobic or anaerobic fumarate-reducing conditions (Fig. 5a). By contrast, no inhibition of anaerobic growth was observed in EC*metR*^+^, consistent with the absence of genes related to anaerobic respiration or cytochrome *c* synthesis in the known *E. coli* MetR regulon^14^. Further research is needed to determine whether the regulation of anaerobic electron transfer systems by MetR is conserved within a taxonomic group that includes both *Shewanella* and *Aeromonas* (e.g., the VAAP subgroup of Gammaproteobacteria^25^) or if this regulatory mechanism is specific to bacteria with EET pathways that require fine-tuned regulation of cytochrome *c* synthesis.

In conclusion, our results suggest that in certain environments, amino acid sources not only provide the nutrients necessary for protein synthesis, but also influence the regulation of bacterial catabolic activity at the transcriptional level. This finding is relevant for the efficient operation of METs using isolates of *Shewanella* and its close relatives cultured in defined media, because the composition and concentration of amino acids in the medium could affect current generation in bioelectrochemical reactors through the regulation of EET activity. However, our results do not exclude the possibility that regulatory factors other than CRP and MetR contribute to the expression of EET-related genes in MR-1. Further investigation is needed to fully understand the molecular mechanisms by which this strain regulates EET activity. We believe that future studies will elucidate the multiple regulatory mechanisms and environmental factors that govern the catabolic activity of this model bacterium.

## Materials and Methods

### Bacterial strains and culture conditions

The bacterial strains used in this study are listed in Supplementary Table S1. *S. oneidensis* and *A. hydrophila* strains were cultured at 30°C in LB medium or minimal medium^11^ containing 10 mM lactate as the carbon and energy source (LMM). *E. coli* was cultured at 37°C in LB medium or M9 minimal medium^35^ containing 20 mM glycerol as the carbon and energy source. Glycerol was used as substrate to inhibit fermentative growth in the absence of external electron acceptors. For aerobic culture, 5 mL of LB, LMM, or M9 glycerol medium in a test tube (30 mL capacity) was inoculated with a bacterial strain at an initial optical density at 600 nm (OD_600_) of 0.05 and was shaken at 180 rpm. For anaerobic culture, 5 or 80 mL of medium in a screw-top test tube (10 mL capacity; for growth curve analysis) or vial (100 mL capacity; for transcriptome analysis), respectively, was supplemented with 20 mM TMAO, fumarate, DMSO, or nitrate as the electron acceptor and inoculated with a bacterial strain at an initial OD_600_ of 0.01. A test tube or vial containing the inoculated medium was sealed with a butyl rubber septum, purged with high-purity nitrogen (99.99%), and incubated without shaking. OD_600_ was measured using a mini photo 518R photometer (Taitec, Saitama, Japan). The *E. coli* mating strain (WM6026) required 100 µg/mL 2,6-diaminopimelic acid for growth. When necessary, 15 μg/mL gentamicin was added to the culture medium. The agar plates contained 1.6% Bacto Agar (Becton, Dickinson and Company).

### Operation of EC

A small single-chamber three-electrode EC with a capacity of 18 mL^12^ was used to monitor the electric current generated by *S. oneidensis* strains. The EC was equipped with a graphite felt working electrode (2.25 cm^2^), an Ag/AgCl reference electrode (+0.199 V vs. SHE) (HX-R5, Hokuto Denko, Tokyo, Japan), and a titanium mesh counter electrode (4 cm^2^, φ0.1 mm, 100 mesh/inch; Nilaco, Tokyo, Japan). The EC was filled with 15 mL of LMM supplemented with 170 mM NaCl as electrolyte and inoculated with bacterial cells at an initial OD_600_ of 0.01. Current was monitored using a multichannel potentiostat (VMP3; Biologic, Claix, France), and current density (A/cm^2^) was calculated based on the projected area of the working electrode. Stock solutions of Met, hCys, and 19 other amino acids were prepared at a concentration of 130 mM and individually added to LMM to achieve a final concentration of 0–130 µM.

### Mutant construction

In-frame disruption of *metR* (SO_0817) in MR-1 was performed using a two-step homologous recombination method with the suicide plasmid pSMV10, as described^36,37^. In brief, a 1.6-kb fusion product, consisting of upstream and downstream sequences of *metR* joined by an 18-bp linker sequence, was constructed by PCR and *in vitro* extension using total DNA from MR-1 and primers (listed in Supplementary Table S2). The amplified fusion product was ligated into the SpeI site of pSMV10. The resulting plasmid, pSMV-metR, was introduced into MR-1 by filter mating with *E. coli* WM6026, and transconjugants were screened as described^36,37^, to isolate double-crossover mutants. The disruption of the target gene in the resulting strains was confirmed by PCR. A representative mutant strain in which *metR* was deleted was selected and designated Δ*metR*.

To construct the plasmid pBBR-SOmetR, expressing *metR* of MR-1 from P_lac_, the insert DNA fragment was PCR-amplified from the total DNA of MR-1 using the primers SOmetR-F-EcoRI and SOmetR-R-SpeI (Supplementary Table S2). The amplified product was digested with EcoRI and SpeI and cloned into the corresponding sites of pBBR1MCS-5^38^. The resulting plasmid, pBBR-SOmetR, was introduced into wild-type MR-1 and Δ*metR* by filter mating with *E. coli* WM6026, and the transformants were designated MR-1*metR*^+^ and Δ*metR*-C, respectively.

To construct the plasmids pBBR-AHmetR and pBBR-ECmetR, expressing *metR* of *A. hydrophila* ATCC 7966 and *E. coli* DH5α, respectively, the insert DNA fragments were PCR-amplified from the total DNA of these strains using primers (listed in Supplementary Table S2). Each PCR product was digested with EcoRI and BamHI (for *A. hydrophila metR*) or EcoRI and PstI (for *E. coli metR*) and cloned into the corresponding sites of pBBR1MCS-5 to yield pBBR-AHmetR and pBBR-ECmetR. Each plasmid was introduced into wild-type *A. hydrophila* or *E. coli* DH5α by filter mating with *E. coli* WM6026 or chemical transformation, respectively. *A. hydrophila* harboring pBBR-AHmetR and *E. coli* DH5α harboring pBBR-ECmetR were designated AH*metR*+ and EC*metR*^+^, respectively.

### RNA extraction

For the transcriptomic profiling of MR-1 under Met-supplemented and unsupplemented conditions, cells (*n* = 3 biological replicates) were cultured aerobically in LMM supplemented with or without 130 µM Met and harvested at the logarithmic growth phase (OD_600_ of 0.2–0.3). For the transcriptomic profiling of MR-1*metR*^+^ and MR-1vc, cells (*n* = 4 biological replicates) were cultured anaerobically in LMM containing 20 mM TMAO as the electron acceptor and harvested at the logarithmic growth phase (OD_600_ of 0.13–0.15). Total RNA was extracted from the cells using Trizol reagent (Thermo Fisher Scientific), followed by purification using RNeasy Mini Kit and RNase-free DNase Set (Qiagen, Valencia, CA), according to the manufacturer’s instructions. The quality of the purified RNA was evaluated using Agilent 2100 Bioanalyzer with RNA 6000 Pico reagents and RNA Pico Chips (Agilent Technologies, Santa Clara, CA), according to the manufacturer’s instructions.

### Transcriptome analyses

Transcriptome analysis was performed using a custom DNA microarray for MR-1 (8 × 15K; Agilent Technologies), which was designed^39^ and validated^12,40–42^ in other studies. Cyanine 3 (Cy3)-labeled complementary RNA was synthesized from 50 ng of total RNA using Low Input Quick Amp WT Labeling Kit (Agilent Technologies) and subjected to microarray hybridization, as described^12^. Gene expression data (*n* = 3–4 biological replicates) were normalized and statistically analyzed using the limma software package (version 3.36.2) for R^43^. A paired Student’s *t* test followed by the Benjamini–Hochberg false discovery rate correction was used for statistical analyses. Differential expression for each probe was considered statistically significant when the absolute value of log_2_ fold change (|log_2_ FC|) was >1.0 at *p* <0.05.

### Quantitative RT-PCR

Quantitative RT-PCR was performed using a LightCycler 1.5 instrument (Roche, Indianapolis, IN, USA), as described^44^. Briefly, the PCR reaction mixture contained 15 ng of total RNA, 1.3 µL of 50 mM Mn(OAc)_2_ solution, 7.5 µL of LightCycler RNA Master SYBR Green I (Roche), and 0.15 µM primers (listed in Supplementary Table S2). To generate the standard curves, the DNA fragments of the target genes were amplified by PCR using Ex Taq DNA polymerase (Takara, Tokyo, Japan) and primer sets (listed in Supplementary Table S2). These were purified by gel electrophoresis using QIAEX II Gel Extraction Kit (Qiagen), according to the manufacturer’s instructions. Standard curves were generated by amplifying a dilution series of the purified DNA fragments of each gene. The specificity of quantitative PCR was verified by dissociation-curve analysis. The expression levels of the target genes were normalized to the expression level of the 16S rRNA gene.

### Purification of MetR

To construct a plasmid expressing MetR with a histidine tag at the C-terminal (C-his-MetR), *metR* was PCR-amplified from the total DNA of MR-1 using the primers metR-F-NdeI and metR-R-XhoI (Supplementary Table S2). The PCR product was digested with NdeI and XhoI and cloned between the corresponding sites of pET-26b(+) (Merck, Darmstadt, Germany). The resulting plasmid, pET-metR (Supplementary Table 1), was introduced into *E. coli* BL21 (DE3). The transformed cells were cultured at 30°C in 300-mL baffled Erlenmeyer flasks containing 100 mL 2× yeast extract-tryptone medium supplemented with Km. Isopropyl-1-thio-β-D-galactopyranoside (final concentration 0.5 mM) was added when the OD_600_ reached 0.3. After overnight culture at 16°C, the cells were harvested by centrifugation, and C-his-MetR was extracted via ultrasonication and purified using QuickPick IMAC Metal Affinity Kit for Proteins (Bio-Nobile, Turku, Finland), as described^45^. The purified protein samples were analyzed by SDS-PAGE and immediately used for subsequent experiments.

### EMSA

EMSA was performed as described^46^ with some modifications. Cy3-labeled 40-bp DNA probes were generated by annealing the complementary single-strand oligonucleotides (listed in Supplementary Table S2). DNA-binding reactions were performed in 20 μL of reaction mixture containing 50 mM Tris-HCl (pH 8.0), 0.5 mM EDTA (pH 8.0), 2 mM MgSO_4_, 1 mM dithiothreitol, 100 µg/mL bovine serum albumin, 10 μg/mL poly(dI-dC) (Sigma-Aldrich), 4 mM hCys, 20% (v/v) glycerol, 1 nM Cy3-labeled DNA probe, and 0–500 ng of C-his-MetR. The mixture was incubated at 20°C for 10 min and loaded onto a nondenaturing 5%–20% gradient polyacrylamide gel (E-T520L, ATTO, Tokyo, Japan). Electrophoresis was performed at 200 V at 4°C in Tris-borate-EDTA buffer for 40 min. Fluorescence gel images were obtained using Typhoon FLA 9000 (GE Healthcare, Milwaukee, WI, USA).

### *In silico* prediction of regulatory sequences

The PSSM for MetR-binding sequences in MR-1 was constructed from 13-bp motifs identified by EMSA using the PSSM-convert program^47^. The PSSM for the CRP-binding sequences was obtained from the RegulonDB database^48^. To identify MetR-or CRP-binding sites, 500-bp upstream sequences of the individual genes in MR-1 or *A. hydrophila* were retrieved and scanned with each PSSM using the matrix-scan function of Regulatory Sequence Analysis Tools^49^ with default parameters. Sequence logos were generated to visualize the PSSMs using WebLogo^50^. A sequence logo for the *E. coli* MetR-binding motif was generated based on 11 known MetR-binding sequences in *E. coli* K-12 obtained from the RegPrecise database^51^.

## Statistical analysis

Data were statistically evaluated using Student’s *t* test or one-way analysis of variance (ANOVA) followed by Tukey’s honestly significant difference (HSD) test using JMP Pro 14.1.0 software (SAS Institute, Cary, NC, USA). Differences were considered statistically significant at a *P* value of <0.05.

## Data Availability

The authors declare that all data supporting the findings of this study are available within the article and its Supplementary Information Files or from the corresponding author upon reasonable request. The microarray data obtained in this study have been deposited in the NCBI Gene Expression Omnibus (GEO) under the accession numbers GSE282263 and GSE282264.

## Acknowledgments

This work was supported by JSPS KAKENHI Grant Numbers 18K05399, 21H02111, and 24K01672.

## Author Contributions

H.M. conducted the majority of the experimental work; K.T. contributed to the quantitative RT-PCR analysis; A.H. contributed to the measurements of current generation in ECs; E.Y. contributed to the transcriptome analysis; T.K. contributed to the EMSA experiments; A.K. conceived and designed the study; K.W. supervised the study; and A.K. and K.W. wrote the manuscript. All authors discussed the results and provided feedback on the manuscript.

## References

1. Russell J. B., Cook G. M. Energetics of bacterial growth: balance of anabolic and catabolic reactions. Microbiol Rev 59, 48–62 (1995).

2. Decker K., Jungermann K., Thauer R. K. Energy production in anaerobic organisms. Angew Chem Int Ed Engl 9, 138–158 (1970).

3. Chubukov V., Gerosa L., Kochanowski K., Sauer U. Coordination of microbial metabolism. Nat Rev Microbiol 12, 327–340 (2014).

4. Venkateswaran K., et al. Polyphasic taxonomy of the genus *Shewanella* and description of *Shewanella oneidensis* sp. nov. Int J Syst Bacteriol 49 **Pt** **2**, 705–724 (1999).

5. Nealson K. H., Saffarini D. Iron and manganese in anaerobic respiration: environmental significance, physiology, and regulation. Annu Rev Microbiol 48, 311–343 (1994).

6. Fredrickson J. K., et al. Towards environmental systems biology of *Shewanella*. Nat Rev Microbiol 6, 592–603 (2008).

7. Kouzuma A. Molecular mechanisms regulating the catabolic and electrochemical activities of *Shewanella oneidensis* MR-1. Biosci Biotechnol Biochem 85, 1572–1581 (2021).

8. Ikeda S., et al. *Shewanella oneidensis* MR-1 as a bacterial platform for electro-biotechnology. Essays Biochem 65, 355–364 (2021).

9. Saffarini D. A., Schultz R., Beliaev A. Involvement of cyclic AMP (cAMP) and cAMP receptor protein in anaerobic respiration of *Shewanella oneidensis*. J Bacteriol 185, 3668–3671 (2003).

10. Kasai T., Kouzuma A., Nojiri H., Watanabe K. Transcriptional mechanisms for differential expression of outer membrane cytochrome genes *omcA* and *mtrC* in *Shewanella oneidensis* MR-1. BMC Microbiol 15, 68 (2015).

11. Ikeda S., Tomita K., Nakagawa G., Kouzuma A., Watanabe K. Supplementation with amino acid sources facilitates fermentative growth of *Shewanella oneidensis* MR-1 in defined media. Appl Environ Microbiol, e0086823 (2023).

12. Hirose A., Kasai T., Aoki M., Umemura T., Watanabe K., Kouzuma A. Electrochemically active bacteria sense electrode potentials for regulating catabolic pathways. Nat Commun 9, 1083 (2018).

13. Hondorp E. R., Matthews R. G. Methionine. EcoSal Plus 2, (2006).

14. Leyn S. A., et al. Comparative genomics of transcriptional regulation of methionine metabolism in *Proteobacteria*. PLoS One 9, e113714 (2014).

15. Cordova C. D., Schicklberger M. F., Yu Y., Spormann A. M. Partial functional replacement of CymA by SirCD in *Shewanella oneidensis* MR-1. J Bacteriol 193, 2312–2321 (2011).

16. Dailey H. A., et al. Prokaryotic heme biosynthesis: multiple pathways to a common essential product. Microbiol Mol Biol Rev 81, (2017).

17. Jin M., et al. Unique organizational and functional features of the cytochrome *c* maturation system in *Shewanella oneidensis*. PLoS One 8, e75610 (2013).

18. Weissbach H., Brot N. Regulation of methionine synthesis in *Escherichia coli*. Mol Microbiol 5, 1593–1597 (1991).

19. Urbanowski M. L., Stauffer G. V. Role of homocysteine in *metR*-mediated activation of the *metE* and *metH* genes in *Salmonella typhimurium* and *Escherichia coli*. J Bacteriol 171, 3277–3281 (1989).

20. Urbanowski M. L., Stauffer G. V. Genetic and biochemical analysis of the MetR activator-binding site in the *metE metR* control region of *Salmonella typhimurium*. J Bacteriol 171, 5620–5629 (1989).

21. Chatterjee J., Miyamoto C. M., Zouzoulas A., Lang B. F., Skouris N., Meighen E. A. MetR and CRP bind to the *Vibrio harveyi* lux promoters and regulate luminescence. Mol Microbiol 46, 101–111 (2002).

22. Dos Santos J. P., Iobbi-Nivol C., Couillault C., Giordano G., Méjean V. Molecular analysis of the trimethylamine N-oxide (TMAO) reductase respiratory system from a *Shewanella* species. J Mol Biol 284, 421–433 (1998).

23. Lemaire O. N., Honore F. A., Jourlin-Castelli C., Mejean V., Fons M., Iobbi-Nivol C. Efficient respiration on TMAO requires TorD and TorE auxiliary proteins in *Shewanella oneidensis*. Res Microbiol 167, 630–637 (2016).

24. Conley B. E., Intile P. J., Bond D. R., Gralnick J. A. Divergent Nrf family proteins and mtrcab homologs facilitate extracellular electron transfer in *Aeromonas hydrophila*. Appl Environ Microbiol 84, 1–14 (2018).

25. Williams K. P., et al. Phylogeny of gammaproteobacteria. J Bacteriol 192, 2305–2314 (2010).

26. Augustus A. M., Spicer L. D. The MetJ regulon in gammaproteobacteria determined by comparative genomics methods. BMC Genomics 12, 558 (2011).

27. Urbanowski M. L., Stauffer L. T., Plamann L. S., Stauffer G. V. A new methionine locus, metR, that encodes a trans-acting protein required for activation of metE and metH in Escherichia coli and Salmonella typhimurium. J Bacteriol 169, 1391–1397 (1987).

28. Raiford D. W., Heizer E. M., Jr., Miller R. V., Akashi H., Raymer M. L., Krane D. E. Do amino acid biosynthetic costs constrain protein evolution in *Saccharomyces cerevisiae*? J Mol Evol 67, 621–630 (2008).

29. Ross D. E., Brantley S. L., Tien M. Kinetic characterization of OmcA and MtrC, terminal reductases involved in respiratory electron transfer for dissimilatory iron reduction in *Shewanella oneidensis* MR-1. Appl Environ Microbiol 75, 5218–5226 (2009).

30. Sturm G., Richter K., Doetsch A., Heide H., Louro R. O., Gescher J. A dynamic periplasmic electron transfer network enables respiratory flexibility beyond a thermodynamic regulatory regime. ISME J 9, 1802–1811 (2015).

31. Bogard R. W., Davies B. W., Mekalanos J. J. MetR-regulated *Vibrio cholerae* metabolism is required for virulence. mBio 3, 1–8 (2012).

32. Pan X., et al. LysR-Type Transcriptional regulator MetR controls prodigiosin production, methionine biosynthesis, cell motility, H2O2 tolerance, heat tolerance, and exopolysaccharide synthesis in Serratia marcescens. Appl Environ Microbiol 86, (2020).

33. Yeung A. T., et al. Swarming of *Pseudomonas aeruginosa* is controlled by a broad spectrum of transcriptional regulators, including MetR. J Bacteriol 191, 5592–5602 (2009).

34. Barchinger S. E., et al. Regulation of Gene expression in *Shewanella oneidensis* MR-1 during electron acceptor limitation and bacterial nanowire formation. Appl Environ Microbiol 82, 5428–5443 (2016).

35. Miller J. Experiments in molecular genetics. CSH NY, (1972).

36. Matsumoto A., Koga R., Kanaly R. A., Kouzuma A., Watanabe K. Identification of a diguanylate cyclase that facilitates biofilm formation on electrodes by *Shewanella oneidensis* MR-1. Appl Environ Microbiol 87, (2021).

37. Saltikov C. W., Newman D. K. Genetic identification of a respiratory arsenate reductase. Proc Natl Acad Sci U S A 100, 10983–10988 (2003).

38. Kovach M. E., et al. Four new derivatives of the broad-host-range cloning vector pBBR1MCS, carrying different antibiotic-resistance cassettes. Gene 166, 175–176 (1995).

39. Kouzuma A., Oba H., Tajima N., Hashimoto K., Watanabe K. Electrochemical selection and characterization of a high current-generating *Shewanella oneidensis* mutant with altered cell-surface morphology and biofilm-related gene expression. BMC Microbiol 14, 190 (2014).

40. Kasai T., Kouzuma A., Watanabe K. CpdA is involved in amino acid metabolism in *Shewanella oneidensis* MR-1. Biosci Biotechnol Biochem 82, 166–172 (2018).

41. Kasai T., Tomioka Y., Kouzuma A., Watanabe K. Overexpression of the adenylate cyclase gene cyaC facilitates current generation by *Shewanella oneidensis* in bioelectrochemical systems. Bioelectrochemistry 129, 100–105 (2019).

42. Koga R., Matsumoto A., Kouzuma A., Watanabe K. Identification of an extracytoplasmic function sigma factor that facilitates c-type cytochrome maturation and current generation under electrolyte-flow conditions in *Shewanella oneidensis* MR-1. Environ Microbiol 22, 3671–3684 (2020).

43. Smyth G. K. Linear models and empirical bayes methods for assessing differential expression in microarray experiments. Stat Appl Genet Mol Biol 3, Article3 (2004).

44. Kouzuma A., Hashimoto K., Watanabe K. Roles of siderophore in manganese-oxide reduction by *Shewanella oneidensis* MR-1. FEMS Microbiol Lett 326, 91–98 (2012).

45. Kouzuma A., Endoh T., Omori T., Nojiri H., Yamane H., Habe H. Transcription factors CysB and SfnR constitute the hierarchical regulatory system for the sulfate starvation response in *Pseudomonas putida*. J Bacteriol 190, 4521–4531 (2008).

46. Sperandio B., et al. Control of methionine synthesis and uptake by MetR and homocysteine in Streptococcus mutans. Journal of Bacteriology 189, 7032–7044 (2007).

47. Klucar L., Stano M., Hajduk M. phiSITE: database of gene regulation in bacteriophages. Nucleic Acids Res 38, D366–370 (2010).

48. Salgado H., et al. RegulonDB v12.0: a comprehensive resource of transcriptional regulation in *E. coli* K-12. Nucleic Acids Res 52, D255–d264 (2024).

49. Turatsinze J. V., Thomas-Chollier M., Defrance M., van Helden J. Using RSAT to scan genome sequences for transcription factor binding sites and cis-regulatory modules. Nat Protoc 3, 1578–1588 (2008).

50. Crooks G. E., Hon G., Chandonia J. M., Brenner S. E. WebLogo: a sequence logo generator. Genome Res 14, 1188–1190 (2004).

51. Novichkov P. S., et al. RegPrecise 3.0 – A resource for genome-scale exploration of transcriptional regulation in bacteria. BMC Genomics 14, 745 (2013).

