## Supplementary Fig. S1 for "Cross-regulation of amino acid synthesis and catabolic electron transfer in bacteria"

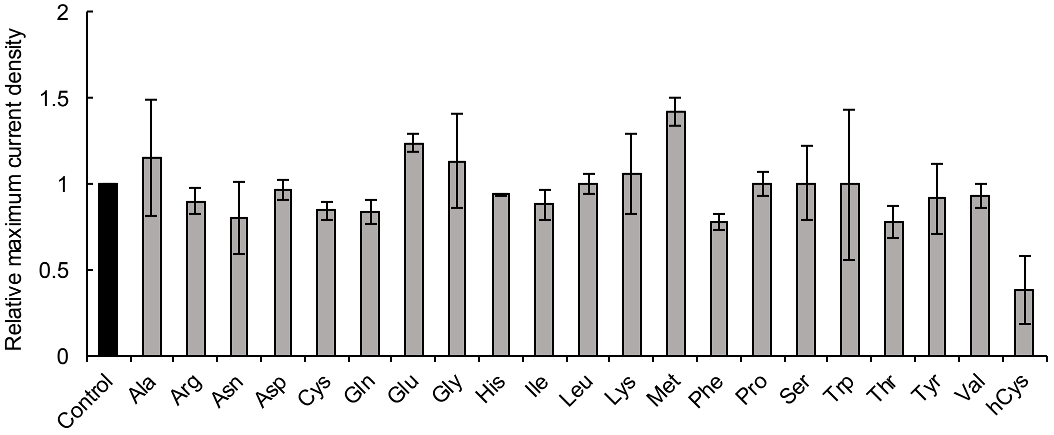


Fig. S1. Comparison of relative current generation by MR-1 in lactate-fed ECs supplemented with one of 20 amino acids and homocysteine (hCys). Each amino acid was added at a concentration of 130 μM. Bars and error bars represent means and standard deviations, respectively, calculated from the results of two independent experiments.
