## Supplementary Fig. S2 for "Cross-regulation of amino acid synthesis and catabolic electron transfer in bacteria"

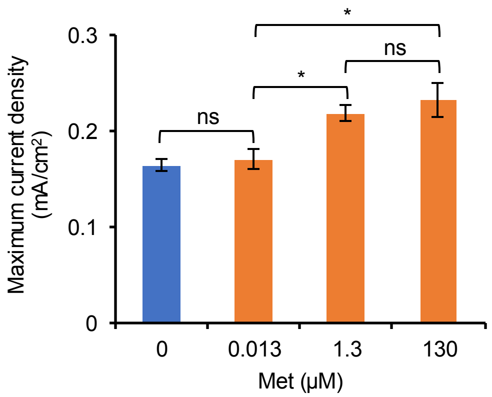


Fig. S2. Maximum current densities generated by MR-1 at different Met concentrations. Bars and error bars indicate means and standard deviations, respectively ($n$ = 3 biological replicates). Asterisks indicate statistically significant differences ($P < 0.05$; one-way ANOVA followed by HSD test); ns, not significant.
