## Supplementary Fig. S3 for "Cross-regulation of amino acid synthesis and catabolic electron transfer in bacteria"

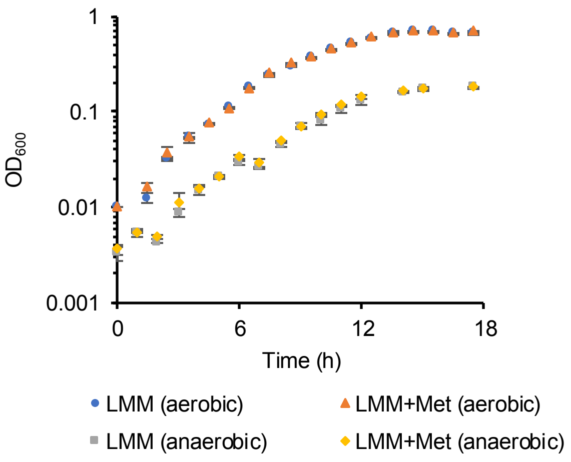


Fig. S3. Growth of MR-1 in LMM supplemented with or without 130 μM Met. Cells were cultured under aerobic or anaerobic (fumarate-reducing) conditions. Data points and error bars represent means and standard deviations, respectively, calculated from the results of two independent experiments.
