## Supplementary Fig. S4 for "Cross-regulation of amino acid synthesis and catabolic electron transfer in bacteria"

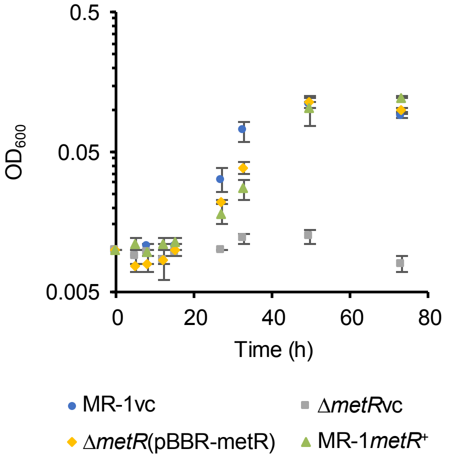


Fig. S4. Growth of MR-1vc, Δ*metR*vc, Δ*metR*(pBBR-metR), and MR-1*metR*$^{+}$ during anaerobic cultivation in LMM supplemented with TMAO as the electron acceptor. Data points and error bars represent means and standard deviations, respectively, calculated from the results of two independent experiments.
