## Supplementary Fig. S5 for "Cross-regulation of amino acid synthesis and catabolic electron transfer in bacteria"

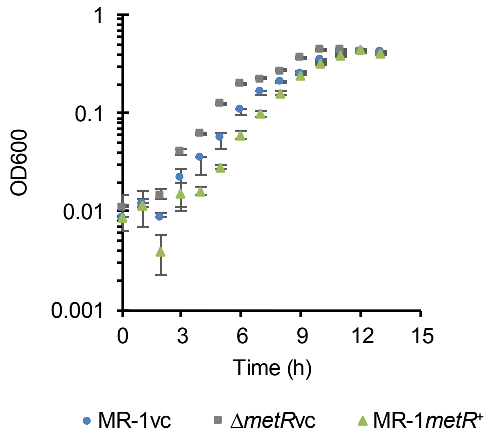


Fig. S5. Growth of Δ*metR*vc during anaerobic cultivation in LB supplemented with TMAO as the electron acceptor. For comparison, the growth curves of MR-1vc and MR-1*metR*$^{+}$ under the same culture conditions (shown in Fig. 2) are plotted. Data points and error bars represent means and standard deviations, respectively, calculated from the results of three independent experiments.
