## Supplementary Fig. S6 for "Cross-regulation of amino acid synthesis and catabolic electron transfer in bacteria"

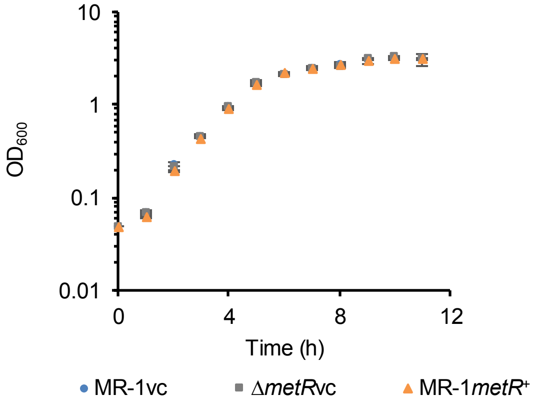


Fig. S6. Growth of MR-1vc, Δ*metR*vc, and MR-1*metR*$^+$ during aerobic cultivation in LB medium. Data points and error bars represent means and standard deviations, respectively, calculated from the results of two independent experiments.
