## Supplementary Fig. S7 for "Cross-regulation of amino acid synthesis and catabolic electron transfer in bacteria"

AAAGAAATAGTCAGATCTCAATCACATTACCCGCTTA

CRP-binding site

AAGTGACTGATAAATACCAATGACTTAGTTAGCCTTA

–35 –10

CAGGT**G**GGATCTAATTCCACACATTAAATACGACTAA

+1

TGAGATTTGTTCTTATTTGATATGTGTTTAATGATG

MetR-binding site *omcA*

Fig. S7. Locations of the CRP- and MetR-binding sites upstream of *omcA*. The transcription start site (+1) of *omcA* is shown in bold, with the putative −35 and −10 promoter sequences underlined. The positions of CRP-binding and transcription start sites are based on other studies (Shao et al., 2014, mBio 5:e01398–14; Kasai et al., 2015, BMC Microbiol 15:68).
