## Supplementary Fig. S8 for "Cross-regulation of amino acid synthesis and catabolic electron transfer in bacteria"

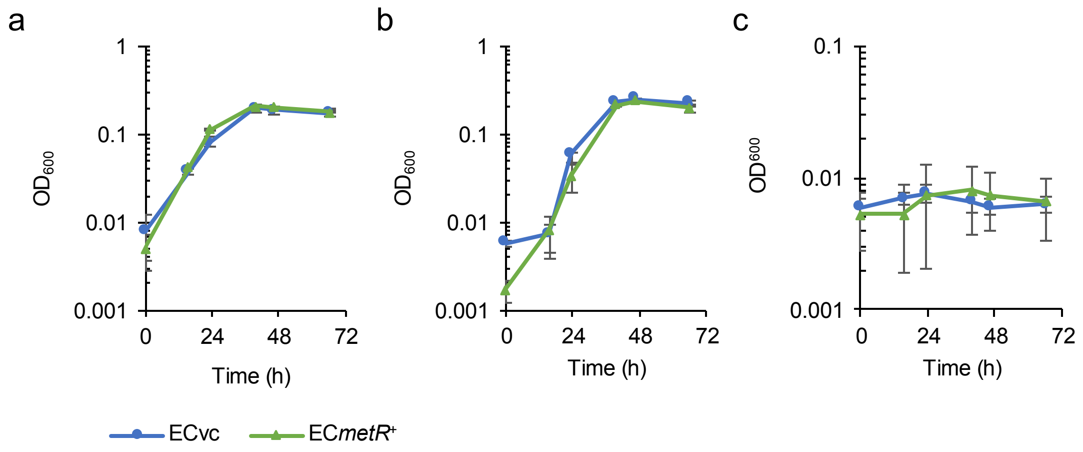


Fig. S8. Anaerobic growth of the ECvc and EC*metR*[+] strains. (a) Growth curves under fumarate-reducing conditions. (b) Growth curves under nitrate-reducing conditions. (c) Growth curves in the absence of electron acceptors as the negative control. In these experiments, cells were cultured in M9 medium supplemented with 20 mM glycerol as a growth substrate to inhibit fermentative growth in the absence of electron acceptors. Data points and error bars represent means and standard deviations, respectively, calculated from the results of three independent experiments.
