## Supplementary Table S1 for "Cross-regulation of amino acid synthesis and catabolic electron transfer in bacteria"

**Supplementary Table S1** Bacterial strains used in this study.

| Strain or plasmid | Relevant characteristic | Source or reference |
|---|---|---|
| **Bacterial strains** | | |
| *Escherichia coli* | | |
| WM6026 | Donor strain for conjugation; *lacI*$^{q}$, *rrnB3*, DE*lacZ4787*, *hsdR514*, DE(*araBAD*)*567*, *E*(*rhaBAD*)*568*, *rph-1*, *att-lambda*::*pAE12*-del(*oriR6K-cat*::*frt5*), DE(*endA*)::*frt*, *uidA*(*delMluI*)::*pir*(wt), *attHK*::pJK1006-*del1/2* (del*oriR6K-cat*::*frt5,* del*trfA*::*frt*) | William Metcalf, University of Illinois |
| DH5α | F$^{-}$, Φ80d*lacZ*ΔM15, Δ(*lacZYA-argF*)U169, *deoR*, recA*1*, *endA*1, *hsdR*17(r$_{K}^{-}$, m$_{K}^{+}$), *phoA*, *supE*44, λ$^{-}$, *thi*-1, *gyrA*96, *relA*1 | Takara |
| ECvc | DH5α harboring pBBR1MCS-5, Gm$^{r}$ | This study |
| EC*metR*$^{+}$ | DH5α harboring pBBR-ECmetR, Gm$^{r}$ | This study |
| *Shewanella oneidensis* | | |
| MR-1 | Wild type | ATCC |
| MR-1$_{vc}$ | MR-1 harboring pBBR1MCS-5, Gm$^{r}$ | This study |
| MR-1*metR*$^{+}$ | MR-1 harboring pBBR-SOmetR, Gm$^{r}$ | This study |
| Δ*metR* | SO_0817 (*metR*) disrupted | This study |
| Δ*metR*-C | Δ*metR* harboring pBBR-SOmetR, Gm$^{r}$ | This study |
| *Aeromonas hydrophila* | | |
| ATCC 7966 | Wild type | ATCC |
| AHvc | ATCC 7966 harboring pBBR1MCS-5, Gm$^{r}$ | This study |
| AH*metR*$^{+}$ | ATCC 7966 harboring pBBR-AHmetR, Gm$^{r}$ | This study |
| **Plasmids** | | |
| pBBR1MCS-5 | Broad-host-range vector, *lacZ* promoter, Gm$^{r}$ | Kovach *et al.* 1995[*] |
| pBBR-SOmetR | pBBR1MCS-5-based plasmid expressing MR-1 *metR* (SO_0817) | This study |
| pBBR-AHmetR | pBBR1MCS-5-based plasmid expressing *A. hydrophila metR* (AHA_1955) | This study |
| pBBR-ECmetR | pBBR1MCS-5-based plasmid expressing *E. coli metR* | This study |
| pSMV10 | 9.1 kb mobilizable suicide vector; *oriR6K*, *mobRP4*, *sacB*, Km$^{r}$, Gm$^{r}$ | Doug Lies, California Institute of Technology |
| pSMV-metR | pSMV10-based plasmid for MR-1 *metR* disruption | This study |
| pET-26b(+) | Expression vector, T7 promoter | Merck |
| pET-metR | pET-26b(+)-based plasmid expressing C-his-*metR* | This study |

[*]Kovach M. E.*, et al.* Four new derivatives of the broad-host-range cloning vector pBBR1MCS, carrying different antibiotic-resistance cassettes. *Gene* **166**, 175-176 (1995).
