## Supplementary Table S2 for "Cross-regulation of amino acid synthesis and catabolic electron transfer in bacteria"

**Supplementary Table S2** Primers used in this study.

| Primer | Sequence (5′–3′) | Modification, for use |
|---|---|---|
| metR-5-O-SpeI | GAAGGTAGACTAGTAATCCTGACCCACGGTTTTG | SpeI, *metR* disruption |
| metR-5-I | CTGACCAACAGCTGACACGGTTCGCAGATGTCTTAG | Linker, *metR* disruption |
| metR-3-I | GTGTCAGCTGTTGGTCAGCGCCAGCAGAAATCTAAC | Linker, *metR* disruption |
| metR-3-O-SpeI | GAAGGTAGACTAGTGAAGAGCAAGTAATTGGCCG | SpeI, *metR* disruption |
| SOmetR-F-EcoRI | CGGAATTCTGAGATAAGCCTATTCGG | EcoRI, pBBR-SOmetR construction |
| SOmetR-R-SpeI | GGACTAGTTAATCCGCTCTAAGAACTGC | SpeI, pBBR-SOmetR construction |
| AHmetR-F-EcoRI | CCGGAATTCGGCACAACAAGGAGGGATACC | EcoRI, pBBR-AHmetR construction |
| AHmetR-R-BamHI | CGCGGATCCTTACTGCGCCTTTACGTCGCC | BamHI, pBBR-AHmetR construction |
| ECmetR-F-EcoRI | CCGGAATTCCGAAGTGAAGGACTTTCATG | EcoRI, pBBR-ECmetR construction |
| ECmetR-R-PstI | AAAACTGCAGTTACAGGCGCGCTGGTGATCC | PstI, pBBR-ECmetR construction |
| qRT-omcA-F | GGATACGGCGTTGAAGATGT | qRT-PCR for *omcA* (SO_1779) |
| qRT-omcA-R | TGGTATCCGTTCCATTCCAT | qRT-PCR for *omcA* (SO_1779) |
| qRT-fccA-F | GTAATTATCGGCTCTGGTGGTG | qRT-PCR for *fccA* (SO_0970) |
| qRT-fccA-R | GCCTGTGGCTTAGTTTCTGC | qRT-PCR for *fccA* (SO_0970) |
| qRT-dmsB-F | GCGCCGTGTTTATGAATATGGC | qRT-PCR for *dsmB* (SO_1430) |
| qRT-dmsB-R | GCAGGCTTTAACACACACAGGC | qRT-PCR for *dsmB* (SO_1430) |
| qRT-metE-F | GAAGGTGTGGGCTTTACCAA | qRT-PCR for *metE* (SO_0818) |
| qRT-metE-R | AATCAACCGTCATGGCTTTC | qRT-PCR for *metE* (SO_0818) |
| qRT-16S-F | AGCGCAACCCCTATCCTTAT | qRT-PCR for MR-1 16S rRNA |
| qRT-16S-R | CGTAAGGGCCATGATGACTT | qRT-PCR for MR-1 16S rRNA |
| metR-F-NdeI | CATATGATCGAACTAAGACATCTGC | NdeI, pET-metR construction |
| metR-R-XhoI | CTCGAGGTTAGATTTCTGCTGGCGAG | XhoI, pET-metR construction |
| omcA-probe-F | CCACACATTAAATACGACTAATGAGATTTGTTCTTATTTG | *omcA* probe for EMSA |
| omcA-probe-R | CAAATAAGAACAAATCTCATTAGTCGTATTTAATGTGTGG | 5'-Cy3, *omcA* probe for EMSA |
| omcAmu-probe-F | CCACACATTAAATACGACTAAGGGGATTTGGGGGTATTTG | *omcA*-m probe for EMSA |
| omcAmu-probe-R | CAAATACCCCCAAATCCCCTTAGTCGTATTTAATGTGTGG | 5'-Cy3, *omcA*-m probe for EMSA |
| cymA-probe-F | GTATTAGCTGGAGTTGAAGTACTCTAACGCTCTGCTAAGC | *cymA* probe for EMSA |
| cymA-probe-R | GCTTAGCAGAGCGTTAGAGTACTTCAACTCCAGCTAATA | 5'-Cy3, *cymA* probe for EMSA |
| glyA-probe-F | CTCAACTCGGCATTGAGGTGCATTCAAGTTAAAACTGTAG | *glyA* probe for EMSA |
| glyA-probe-R | CTACAGTTTTAACTTGAATGCACCTCAATGCCGAGTTGAG | 5'-Cy3, *glyA* probe for EMSA |
| luxS-probe-F | GCATCTGACTGATATGAGATGATTTCATCTTAAGTTATC | *luxS* probe for EMSA |
| luxS-probe-R | GATAACTTAAGATGAAATCATCTCATATCAGTCAGATGC | 5'-Cy3, *luxS* probe for EMSA |
| metR-probe-F | GATTTCAAGCGCAACATGAGCGAAATTCACTTGGAGTGATC | *metR* probe for EMSA |
| metR-probe-R | GATCACTCCAAGTGAATTTCGCTCATGTTGCGCTTGAAATC | 5'-Cy3, *metR* probe for EMSA |
| dmsE-probe-F | CTTATGGTGTTTTTGAGAATGATTTTCTTTTATTTGATTTC | *dmsE* probe for EMSA |
| dmsE-probe-R | GAAATCAAATAAAAGAAAATCATTCTCAAAAACACCATAAG | 5'-Cy3, *dmsE* probe for EMSA |
